# Ladostigil attenuates the oxidative and ER stress in human neuroblast-like SH-SY5Y cells

**DOI:** 10.1101/2021.08.07.455498

**Authors:** Keren Zohar, Elyad Lezmi, Tsiona Eliyahu, Michal Linial

## Abstract

A hallmark of the aging brain is the robust inflammation mediated by microglial activation. Neuroinflammation resulting from the induction of oxidative stress in neurodegenerative diseases and following brain injury. Chronic treatment of aging rats by ladostigil, a compound with antioxidant and anti-inflammatory function, prevented microglial activation and learning deficits. In this study, we investigate the effect of ladostigil on neuronal-like SH-SY5Y cells. We show that SH-SY5Y cells exposed to acute (by H_2_O_2_) or chronic oxidative stress (by Sin1, 3-morpholinosydnonimine) induced apoptotic cell death. However, in the presence of ladostigil, the decline in cell viability and the oxidative levels were partially reversed. RNA-seq analysis showed that chronic oxidation by Sin1 resulted in coordinated suppression of endoplasmic reticulum (ER) quality control and ER stress response gene sets. Chronic oxidative stress impacted ER proteostasis and induced the expression of numerous lncRNAs. Pre-incubation with ladostigil before exposing SH-SY5Y cells to Sin1 induced Clk1 (Cdc2-like kinase 1) which was implicated in psychophysiological stress in mice and Alzheimer disease. Ladostigil also suppressed the expression of Ccpg1 (Cell cycle progression 1) and Synj1 (Synaptojanin 1) that function in ER-autophagy and endocytic pathways. We postulate that ladostigil alleviated cell damage by oxidation and ER stress. Therefore, it may attenuate neurotoxicity and cell death that accompany chronic stress conditions in the aging brain.

## Introduction

A decline in cognitive function is a hallmark of normal brain aging, manifested by changes in brain morphology and cell type composition [1]. The deficit in learning and memory shows a decreased number of neurons, reduction of synaptic sites, and changes in the properties of dendritic spines [2, 3]. At a cellular level, decline in the brain’s higher functions are associated with apoptotic death [4], reduced mitochondrial function [5], proteotoxicity [6, 7], and enhanced autophagy [8]) among others. A phenomenon cell characteristic of the aging brain is a reduced capacity to cope with stress (e.g., oxidative stress) compared to cells from young and healthy adult brains [9]. A failure in antioxidant defense mechanisms in the aging brain [10, 11] and impaired mitochondrial function [12] are associated with memory decline [13]. Once the cellular compensatory mechanisms can no longer cope with the stress [14], pathological processes are induced [15, 16] as observed following brain injury [17] and in neurodegenerative diseases.

Chronic treatment with ladostigil, a compound with antioxidant and anti-inflammatory activity [18], was shown to prevent further loss in learning and memory [19–21]. In addition, ladostigil treatment restored the morphology and activation state of microglia to pretreatment levels [22]. Other molecular changes induced by aging and attenuated by ladostigil include impairment of neurogenesis, synapse formation, calcium and fatty acid homeostasis [21]

In vitro experiments mimic brain inflammation with a simpler and controllable setting. Specifically, by activating primary neonatal microglia with Lipopolysaccharides (LPS) and ATP [23], the cells secreted TNFα, IL-1β, and IL-6 [24]. These pro-inflammatory cytokines contribute to synapse loss and learning deficits [25]. Using in vitro setting, Ladostigil suppressed TNFα and IL-1β release presumably by blocking Nf-κB from translocating to the nuclei [26].

One of the outcomes of microglial activation (as well as that of macrophages, neutrophils, and endothelial cells) is the accumulation of reactive oxygen species (ROS) which include superoxide, singlet oxygen, hydroxyl radical, and hydrogen peroxide (H_2_O_2_). These molecules can damage proteins, DNA, and lipids at the cell membranes, ultimately leading to pathological changes of cell structure and function [27]. Mitochondria are the primary source of ROS in cells. Upon oxidative stress, apoptosis and mitophagy are activated for attenuating any potential damage [28, 29]. In the mammalian brain, neurons are strongly dependent on oxidative metabolism for ATP production, and therefore, are prone to ROS-dependent damage. An imbalance between ROS production and antioxidants occurs in Alzheimer’s disease, Parkinson’s disease, and other types of dementia of the elderly [30].

In this study, we examined the effect of ladostigil on neuronal-like cells. We used human SH-SY5Y cells, a derivative of SK-N-SH neuroblastoma cells. SH-SY5Y cells possess numerous neuronal-like properties (e.g., dopamine-producing enzymes, glutamate to GABA conversion, and an active neuronal-like secretory system) [31]. We monitored the changes in cell response following ladostigil treatment under varying conditions of oxidative stress. We introduced a mild, chronic oxidative stress that mimics the increased age-dependent mitochondrial burden. The beneficial effect of ladostigil on cells’ viability is attributed to its impact on dissipating elevated oxidative levels. Finally, we performed RNA-seq analysis on cells under stress and with ladostigil treatment. Oxidation-induced stress conditions led to lncRNAs induction and suppression of genes that function in folding, proteostasis, and autophagy. We showed that ladostigil acts on the ER stress and membrane dynamics axis. We discuss ladostigil’s capacity to maintain proteostasis by coping with oxidative and ER stress.

## Materials and Methods

### Materials

MTT (3-(4,5-dimethylthiazol-2-yl)-2,5-diphenyltetrazolium bromide) was used for cell viability. All reagents were purchased from Sigma Aldrich (USA) unless otherwise stated. Ladostigil, (6-(N-ethyl, N- methyl) carbamyloxy)-N propargyl-1(R)-aminoindan tartrate, was a gift from Spero BioPharma (Israel). Media products MEM and F12 (ratio 1:1), heat-inactivated fetal calf serum (10% FCS), L-Alanyl-L Glutamine were obtained from Biological Industries (Beit-Haemek, Israel). All tissue culture materials were purchased from Beth-Haemek (Israel). H_2_O_2_ (Sigma, Germany) was diluted in cold water and the solutions were used within 2 hrs. Sin1 (hydrochloride, 3-(4-5 Morpholinyl) sydnone imine hydrochloride, Linsidomine hydrochloride Sigma Lipofectamine-2000 was used as a transfection reagent according to the manufacturer’s instructions.

### SH-SY5Y cell culture

Human neuroblastoma SH-SY5Y cells were obtained from ATCC (American Type Culture Collection, MD, USA) [32]. Cells were cultured in Minimum Essential Media (MEM and F12 ratio 1:1, 4.5 g/l glucose) with 10% fetal calf serum (FCS) and 1:10 L-Alanyl-L-Glutamine. Cell cultures were incubated at 37°C in a humidified atmosphere of 5% CO_2_. Ladostigil was added to the culture medium 2-hrs prior to the activation of oxidative stress to cells. Cells were tested 24 hrs later (i.e., 26 hrs after the addition of ladostigil).

### Cell viability assay

SH-SY5Y cells were cultured at a density of 2×10^4^ cells per well in 96-well plates, in 200 μl of medium. MTT assay used colorimetric measurement for cell viability where viable cells produced a dark blue formazan product. Cells were treated in the absence or presence of ladostigil, at different concentrations. MTT solution in phosphate-buffered saline (PBS, pH 7.2) was prepared at a working stock of 5 mg/ml. After 24 hrs, culture medium was supplemented with 10 μl of concentrated MTT per well. Absorption was determined in an ELISA-reader at *λ* = 535, using reference at 635 nm. Cell viability was expressed as a percentage of untreated cells. Each experimental condition was repeated 8 times.

### Oxidative stress sensor

Cells were transfected with molecular sensors that allow a change in fluorescence signal normalized by GFP [33]. Cyto-roGFP plasmids and Mito-matrix-roGFP are redox-sensitive biosensors. The plasmids were a gift from the laboratory of D. Reichman (Hebrew University) and are identical to the commercial Addgene vectors (MA, USA) [34]. SH-SY5Y cells were cultured in 6- or 12-well plates at 37°C and 5% CO_2_, to reach a 70-80% confluent level. For transfection, a ratio of 1:1 of the roGFP plasmids (Cyto and Mito-roGFP) were added at room temperature to the culture in MEM and F12 (at a cell density of 2×10^4^ per cm^2^). A total of 5.5 μg of the total DNA mixture was used for the 6-wells, respectively. The reagent transfected with DNA was incubated for 24-36 hrs before analysis by flow cytometry (FACS). The 488-nm laser was used for excitation and the 530 nm optical filter was used for fluorescence detection. The 405-nm laser was used for excitation and the 525 nm optical filter was used for fluorescence detection. Dithiothreitol (DTT) was used for the calibration of maximal reduction, while diamide was used to achieve maximal cells oxidation. Propidium Iodide (PI) was used to quantify the percentage of dead cells. **Supplementary Figure S1** shows the results of DTT and diamine calibration along with the naive untreated cells’ redox state.

### Reverse transcription polymerase–chain reaction (RT-PCR)

Total RNA was extracted with Trizol (Life Technologies, MD, USA), and RT was performed using a Ready-To-Go first-strand synthesis kit (Amersham Pharmacia Biotech, UK) according to the manufacturer’s instructions. RNA was reverse transcribed into cDNA (1 μg), and used in the PCR reaction. The PCR conditions consisted of denaturation at 95°C for 2 min and 35 cycles (10 sec at 95°C, 15 sec at 60°C, and 5 min at 72°C for extension). Glyceraldehyde 3-phosphate dehydrogenase (GAPDH), ribosomal L19 of β-actin were used as internal controls in one reaction confirming that the RT-PCR amplicon sizes were distinctive. The PCR products were separated on 1.5%-2% agarose gel and stained with ethidium bromide, followed by densitometry measurement (using ImageJ). **Table 1** lists the forward and reverse primers used for the RT-PCR. We also performed a real-time PCR reaction using β-actin for normalization.

**Table 1.**
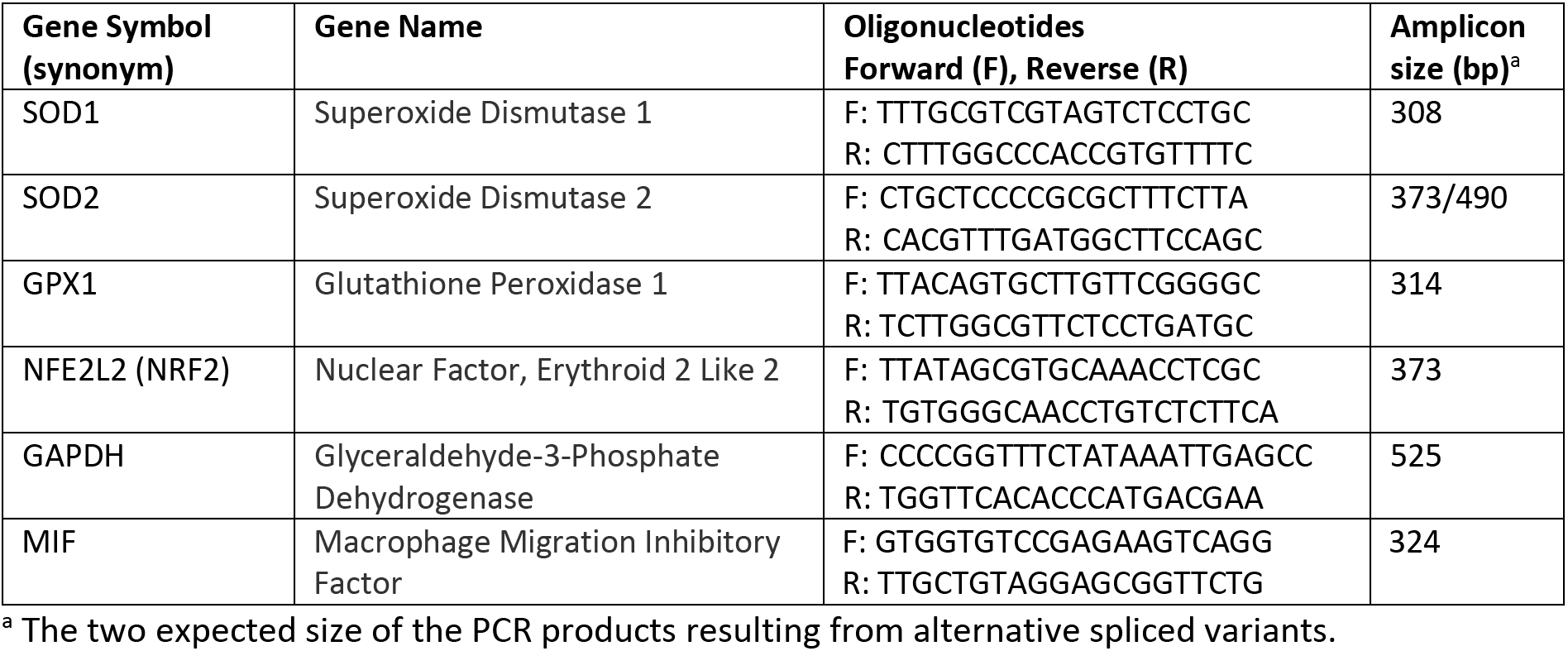
RT-PCR on selected genes participate in cell redox balance.

### RNA sequencing

SH-SY5Y cells were pre-incubated for 2 hrs with ladostigil at a concentration of 5.4 μM. Cells were exposed to Sin1 (t=0) and harvested at 10 hrs or 24 hrs following the addition of ladostigil. Total RNA was extracted using the RNeasy Plus Universal Mini Kit (QIAGEN) according to the manufacturer’s protocol. Total RNA samples (1 μg RNA) were enriched for mRNAs by pull-down of poly(A) and libraries were prepared using the KAPA Stranded mRNA-Seq Kit according to the manufacturer’s protocol. RNA-seq libraries were sequenced using Illumina NextSeq 500 to generate 85 bp single-end reads.

### Differential expression analysis

Next-generation sequencing (NGS) data underwent quality control using FastQC, aligned to the reference genome GRCh38 with STAR aligner [35] using default parameters. Genomic loci were annotated using GENCODE version 37 [36]. Following normalizations, low expressing genes were filtered out by setting a threshold of a minimum of 5 counts-per-million in three samples. Genes with FDR <0.05 and an absolute log fold-change above 0.3 were considered as significantly differentially expressed (unless otherwise mentioned).

### Statistics

All experiments were performed with a minimum of three biological replicates. Principal component analysis was performed using the R-base function “prcomp”. The counts per gene were normalized using the “weighted” Trimmed Mean of M-values (TMM) approach, according to the Bioconductor package EdgeR [37]. Enrichment analyses for pathways used GO pathway based on slim GO biological processes (PANTHER ver 16) [38]). All figures were generated using the ggplot2 R package.

### Data availability

RNA-seq data files were deposited to ArrayExpress under the accession E-MTAB-10817.

## Results

### An immunological signature dominates aged rat brain

Comparing rat brain transcriptomes across several regions from young adults and old rats (6.5 and 22 months, respectively) highlighted many of the age-dependent changes in gene expression [21]. The selected brain regions participate in neuronal plasticity, memory, and information processing. A dominant signature of immune-related genes over-expressed in aged versus young-adult brains. Among these genes are B2m (Beta-2-microglobulin), Cd74 (Major histocompatibility complex, Class II invariant chain), numerous representatives of MHC class II (refer to RT-class 1 and 2 antigens, ~ 13 genes), and more. **Figure 1** shows the expression levels of genes (p-value FDR <0.05) in adults and old rats in three brain regions: frontal cortex (FC), hippocampus (HIP), and perirhinal cortex (PER).

**Figure 1.**
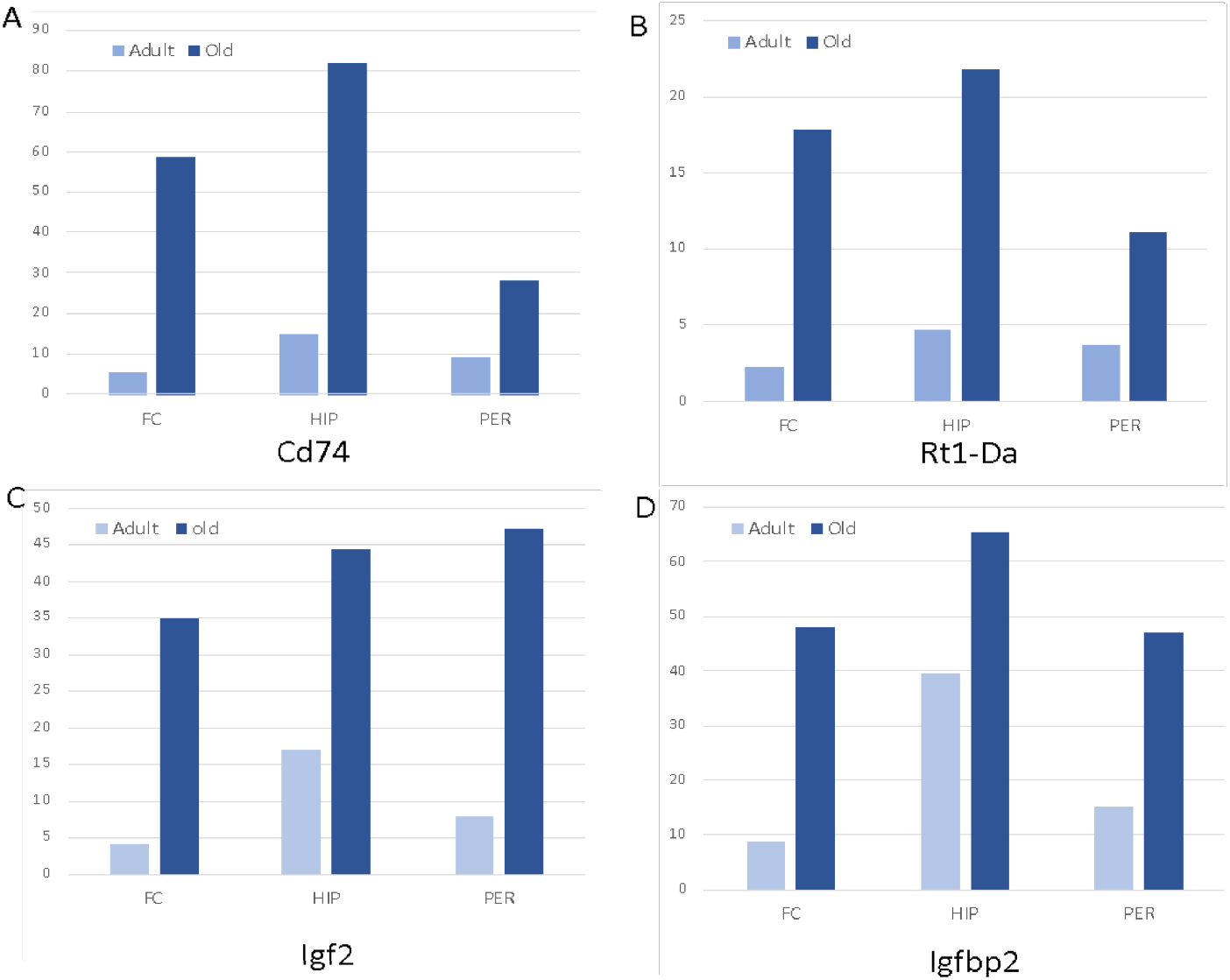
Representative genes that are differentially expressed in rat brains of young adults relative to old animals. Results are based on RNA-seq analysis with three biological samples for each age group and each brain region: the frontal cortex (FC), hippocampus (HIP), and perirhinal cortex (PER). **(A)** The relative expression of Cd74 is 11.0, 5.5, and 3.2-fold in FC, HIP PER, respectively. **(B)** The Rt1-Da, an endogenous antigen-presenting peptide of MHC class II, is strongly elevated in the old brain (8.1, 4.7, and 3.0 folds in FC, HIP PER, respectively). **(C)** The expression of insulin-like growth factor (Igf2) and **(D)** its binding protein (Igfbp2) was increased by 8.3 and 5.5 folds in FC, 2. 6 and 1.6-fold in HIP, and 6.0 and 3.1-fold in PER. The listed genes show a statistical significance of p-value FDR <1E-06.

A coordinated increased expression of Cd74 and Rt1-Da (**Figures 1A–1B**) is attributed to the marked increase in activated microglia in old rat brains. Rt1-Da and Cd74 are specific markers of activated microglia that secrete proinflammatory cytokines (M1 microglia) [39]. A similar increase was observed in insulin-like growth factor 2 (Igf2) and Insulin-like growth factor binding protein 2 (Igfbp2) expression levels (**Figures 1C–1D**). These neurotrophic growth factors act in coordinating neuron-glia cell communication leading to neuronal maturation, remodeling, and plasticity of high-level cognitive functions [40]. Considering the dominant effect of microglia on the RNA-seq profile in the rat brain the direct outcome of ladostigil on neurons was impossible to determine, as subtle effects are likely to be masked by glial cells. To this end, we simplified the experimental setting and looked for a molecular outcome of ladostigil on neuronal-like SH-SY5Y cells under basal and stressed conditions.

### SH-SY5Y cell survival upon acute oxidative stress

In testing the sensitivity of SH-SY5Y cells to a short-lived oxidative insult, we exposed cells to hydrogen peroxide (H_2_O_2_) and measured cell viability using MTT assay (**Figure 2**). Cells that were exposed to a moderate level of H_2_O_2_ (up to 80 μM) remained viable, suggesting that SH-SY5Y cells can dissipate H_2_O_2_ successfully and neutralize its potentially damaging effects (**Figure 2A**).

**Figure 2.**
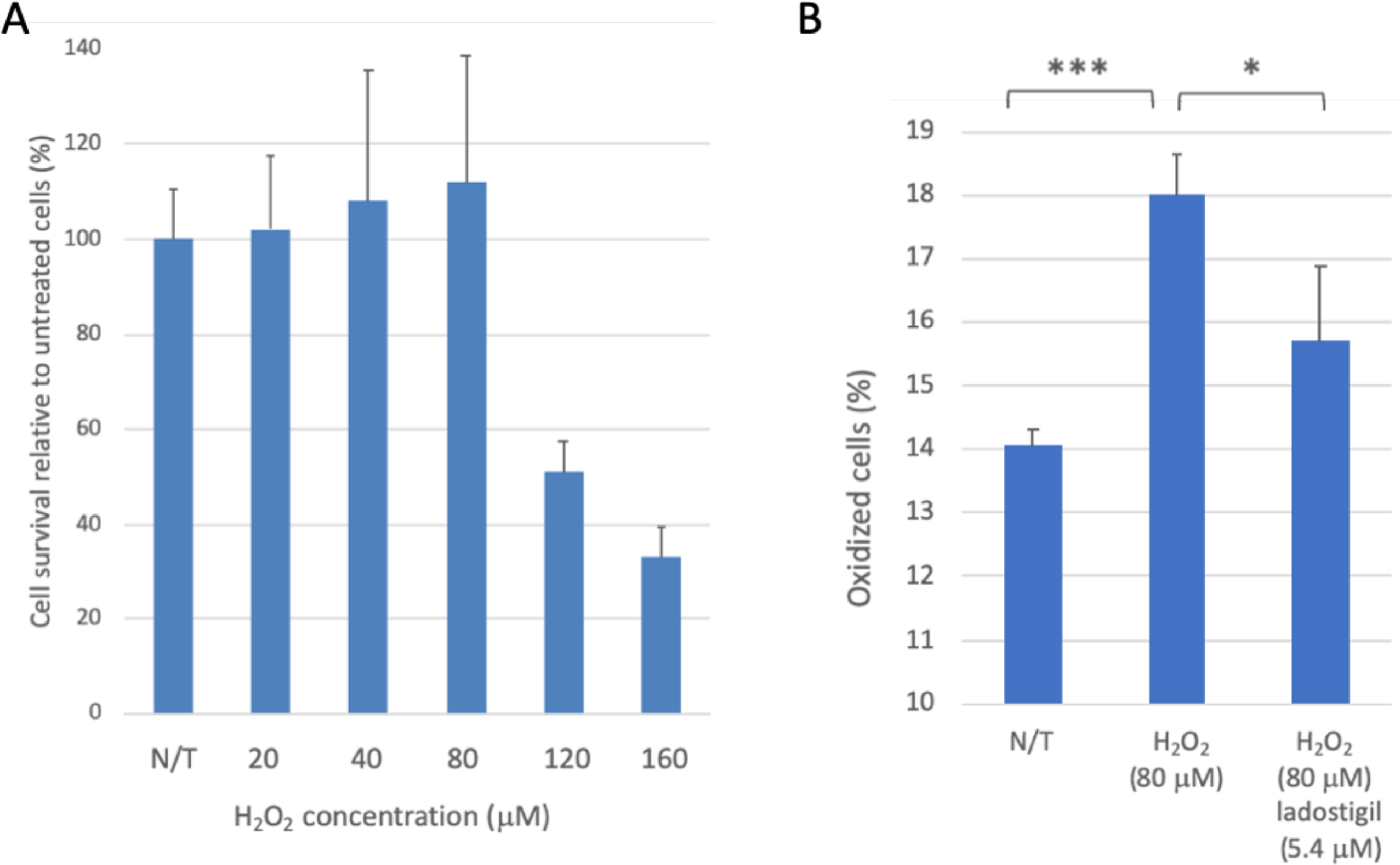
SH-SY5Y cells under acute short-term oxidation stress and ladogtigil treatment. **(A)** Cell viability was analyzed by MTT assay after 24 hrs using increasing concentrations of hydrogen peroxide (H_2_O_2_). **(B)** Cells transfected with plasmid carrying sensors for redox state (see Materials and methods) were analyzed by FACS. In each experiment, the fraction of oxidized cells is determined relative to the untreated cells. The cells were exposed to H_2_O_2_ for 3 hrs with or without pre-incubation with ladostigil (for 1 hr). Each group represents biological triplicates. Statistical test (t-test) showed significance at the level of <0.05 (*) and <1E-03 (***).

To test a possible shift in the cellular redox state in the presence of H_2_O_2_, we utilized oxidation state molecular sensors [41]. Cells were transfected with a mixture of cytosolic and mitochondrial fluorescence sensors for monitoring the redox state (see Materials and Methods, **Supplemental Figure S1**). A marked increase in the fraction of oxidized cells was observed (28%, p-value <1E-03, **Figure 2B**) in cells exposed to H_2_O_2_ (80 μM) for 3 hrs. We found that pre-incubation with ladostigil (for 1 hr, at 5.4 μM) before adding H_2_O_2_, significantly reduced the oxidative state compared to unexposed cells (to 87%, p-value <0.05). Using the same protocol and testing the cell oxidative state 24 hrs after exposure to ladostigil reverted such short-term effects **(Figure 2B)**. We concluded that SH-SY5Y cells cope successfully with short-term oxidation stressors and therefore are less appropriate for measuring the long-term effects of stress on neuronal-like cells. Cells that were pre-incubated for 1 hr before being exposed to the oxidative insult, showed no sign of ladostigil neurotoxicity at varying concentrations (**Supplemental Figure S2**).

### SH-SYS5 redox state under induced long-term oxidation stress

Based on the observations in **Figure 2**, we replaced H2O2 with Sin1 (3-morpholinosydnonimine) [42]. Sin1 is a long-lasting stressor that simultaneously generates nitric oxide (NO) and superoxide (O2−). Notably, microglia (as well as neutrophils, and endothelial cells) produce these reactive molecules under physiological conditions. We calibrated the effect of a mild yet long-term oxidative stress by Sin1 as a substitute for chronic stress occurring along brain’s aging. **Figure 3A** shows the results of cell viability assays in the presence of Sin1. Viability is unaffected and remains stable at a wide range of Sin1 concentrations (50-100 μM). However, a strong reduction in cell viability was recorded at higher concentrations (300 μM; 18% survival). Under such conditions, pre-incubation of ladostigil (1-2 hrs) improved viability to a level of ~75% relative to untreated cells. The cells’ protective effect by ladostigil was not observed at a much higher Sin1 concentration (500 μM, Figure 3A).

**Figure 3.**
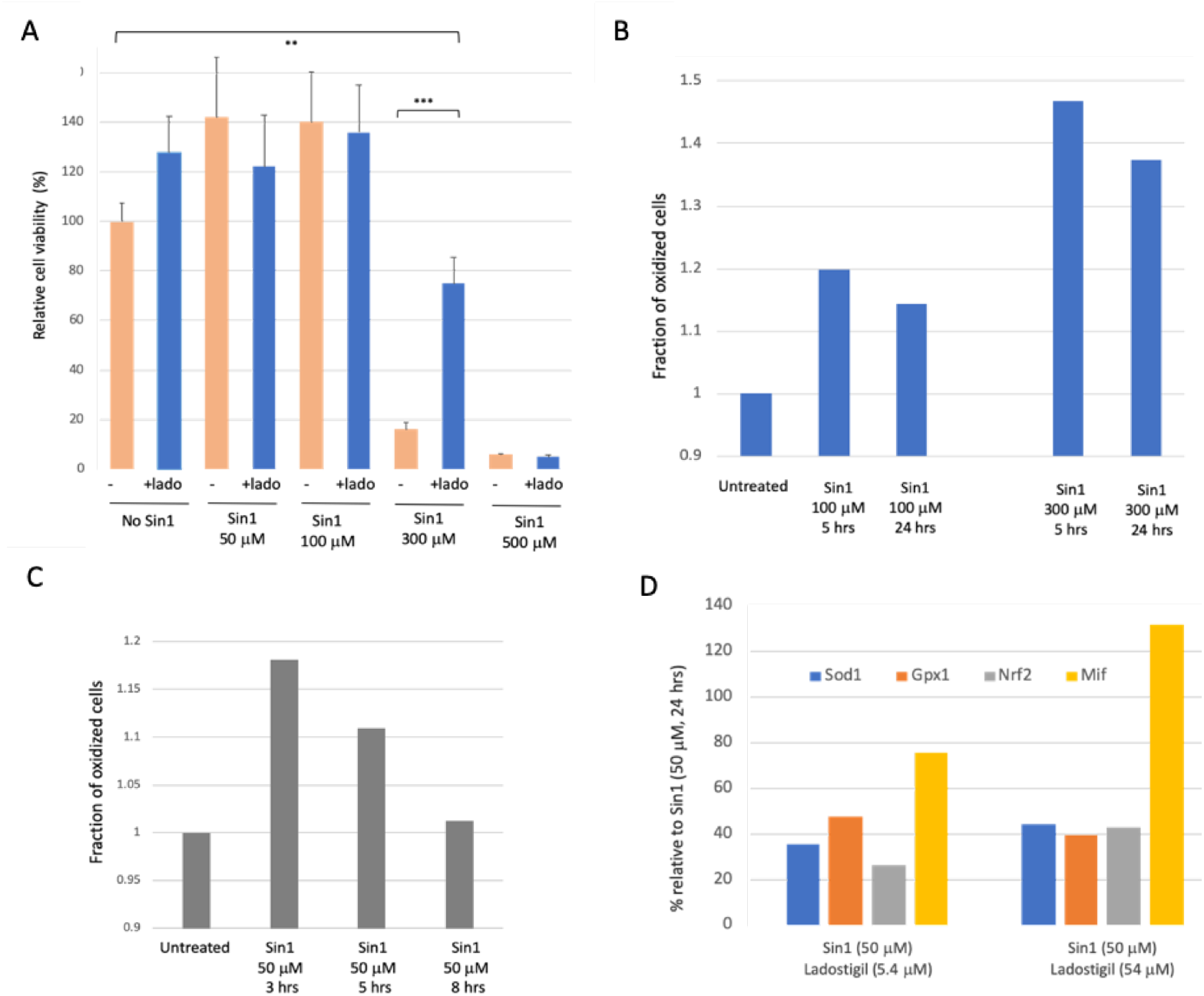
SH-SY5Y cells under chronic long-term oxidation stress and ladogtigil treatment. **(A)** Cell viability upon increasing Sin1 concentration in the presence or absence of ladostigil (5.4 μM). **(B)** Monitoring the redox state of transfected cells of transfected cells express GFP (see Materials and Methods). Cells were exposed to a higher Sin1 concentration (100-300 μM) for 5 and 24 hrs. The fraction of the oxidized cells is determined relative to untreated cells. Each experiment was repeated 4 times with <15% difference from the mean. (**C)** Cells were exposed to low levels of Sin1 (50 μM), and the culture was monitored for its redox status at varying times (3 to 24 hrs). Each experiment was repeated 3 times with below 15% from mean. **(D)** Quantitative real-time PCR of redox-response genes with ladostigil (5.4 μM and 54 μM). Reference is to Sin1 cells without the preincubation of ladostigil (marked as 100%).

We tested whether the effect of Sin1 can be directly monitored by the cell’s redox state, using FACS-based measurements. We combined mitochondrial and cytosolic sensors (at 1:1 ratio, see Materials and Methods) for monitoring the overall cell state. For each experiment, cells were calibrated by applying reducing and oxidizing reagents (DTT and diamide, respectively; **Supplemental Figure S1**). **Figure 3B** shows that at low levels of Sin1 (50 μM), the fraction of oxidized cells was increased (by 18%) but returned to the basal state after 8 hrs. At higher levels of Sin1, an elevated oxidative state lasted 24 hrs, with an increase in the oxidized cell fraction of 14% and 37% for 100 μM and 300 μM of Sin1, respectively (**Figure 3B**). The maximal effect was measured 5 hrs following the insult. We repeated the assay of cells’ redox state while seeking a balance between chronic stress and irreversible cell death (**Figure 3C**).

**Figure 3D** shows the results from real-time quantitative PCR for gene representatives of redox function and ROS-sensitive genes (**Table 1**). We tested the relative expression of selected genes by preincubating ladogtigil before adding Sin1 (2 hrs). The genes tested include the antioxidant superoxide dismutase enzymes Sod1, Glutathione peroxidase 1 (Gp×1), Nfe2l2, a major oxidation sensitive gene regulatory transcription factor (Nrf2) and Mif (Macrophage migration inhibitory factor), a proinflammatory cytokine that is involved in cell death and survival. For Sod1, Gp×1, and Nrf2, ladostigil reduced the expression to ~40% of the maximal induced level by Sin1. An exception is the expression of Mif that was induced by Sin1 but ladostigil further induced its expression.

We extended the analysis to compare enzymes that may protect against apoptotic cell death [43]. **Supplemental Figure S3A** shows the results from RT-PCR gel-assay for Sod1, Sod2, and Gp×1. Sod genes catalyze the dismutation of superoxide radical anion to H2O2 and free oxygen. The cytoplasmic Sod1, only slightly induced by Sin1, while Sod2, a mitochondrial resident enzyme, was induced >2-fold from its basal level. Ladostigil at 5.4 μM reduced Sod2 to the untreated cell’s basal level. A similar pattern was recorded for Glutathione peroxidase 1 (Gp×1) which acts to remove intracellular peroxides. In the presence of ladostigil at a high Sin1 concentration (100 μM) the expression of Gp×1 was maximally suppressed. The results for Sod2, Gp×1 confirm a reduction in the expression of these genes by pre-incubation with ladostigil. Mif was induced by ladostigil in cells stressed by Sin1 (**Supplemental Figure S3B**). This is in accord with Mif’s role as a chaperone, allowing protecting the mitochondria and the ER from accumulated misfolded Sod1 [44]. We concluded that ladostigil takes part in suppressing the elevation in the oxidized state in response to a mild level of Sin1.

### A coordinated suppression of ER chaperones and folding enzymes by a chronic oxidative stress

We performed RNA-seq (in biological triplicates) to compare the transcriptome of untreated cells and Sin1-treated cells. We tested the transcriptome 24 hrs after exposure to monitor the adapted cellular response. Among the genes that are robustly expressed (total 11,532 genes, see Materials and Methods), 84% were unchanged, 9% were induced, and 7% were suppressed by Sin1. Among the unchanged genes, 97% are protein-coding (**Supplemental Table S1**).

**Figure 4A** shows a volcano plot comparing non-treated naïve cells and cells exposed to a mild level of Sin1 (50 μM) after 24 hrs. Note that most statistically significant genes are suppressed by Sin1 (colored blue). We compared the significantly suppressed (**Figures 4B**) and induced (**Figure 4C**) genes following Sin1 treatment relative to non-treated cells and assesses their annotations’ enrichment according to GO (Gene Ontology) biological process terms. The most enriched terms for the suppressed genes (n=402) include different aspects of response to unfolding proteins, de novo folding, and response to ER stress [45]. Testing the enrichment for the induced genes (n=493) resulted in lower enrichment relative to the suppressed genes (**Figures 4B–4C**, x-axis). However, the dominant GO biological process terms concern neuronal development and differentiation. We concluded that Sin1 treatment led to changes in the protein folding response and ER stress, while the induced genes highlight neuronal differentiation.

**Figure 4.**
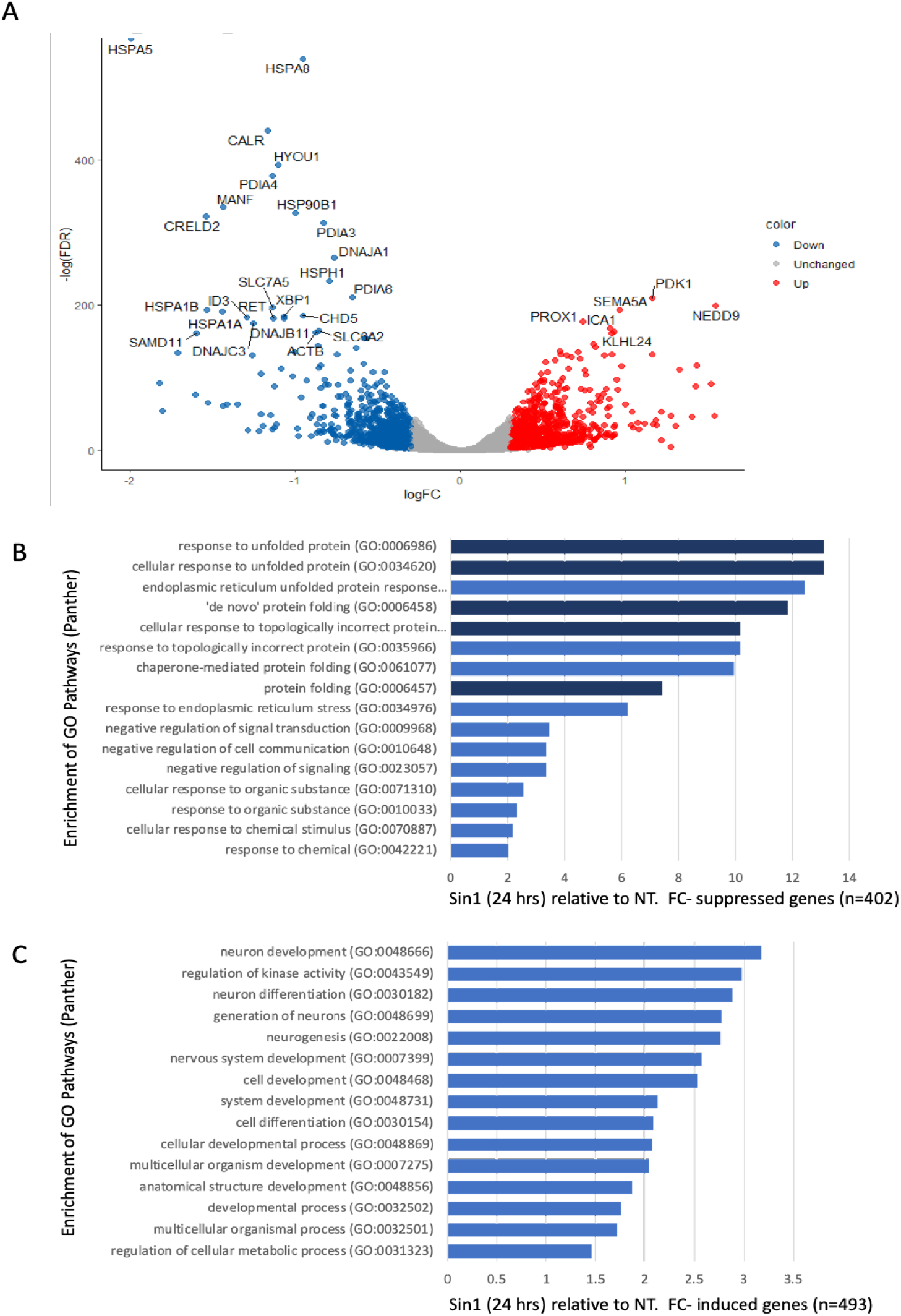
Differential expression and annotation enrichment of SH-SY5Y cells comparing the expression of non-treated naïve cells and cells exposed to Sin1 (50 μM) for 24 hrs**. (A)** Volcano plot of the differential expressed genes that were induced (red) and suppressed (blue) according to the fold change and statistical significance. **(B)** Enrichment of suppressed genes with normalized expression level >5 TMM, and <0.75-fold change. **(C)** Enrichment of induced genes with normalized expression level of >5 TMM and >1.33-fold change. Listed are the terms with >10 genes that are statistically significant (p-value FDR <0.05; blue). Terms with p-value FDR <0.001 are marked by a dark blue. Indicated terms are from the Panther GO process list. FC, fold change. NT, non-treated SH-SY5Y cells.

**Figure 5** summarizes the suppression of all 34 genes which are strongly suppressed (>2 folds; minimal p-value FDR <1E-12). The gene list is enriched with chaperones, folding enzymes form multiprotein complexes of endoplasmic reticulum (ER) function. According to the cellular pathways map (ENRICHR [46]), Dnajc3, Xbp1, Hspa5, c, Hyou1, Calr, Hspa1b, Herpud1, Pdia4, Hsp90b1, Hspa1a are associated with protein processing and ER quality control, showing an enrichment of 40.9 folds (p-value FDR of 2.5E-13). Major chaperones (Hspa5, Hspa1b, Hspa1a, Hsp90b1, Hyou1) belong to the Hsp90 and Hsp70 families, located at the ER lumen. In addition, among the genes suppressed by Sin1 are co-chaperones of BiP (Dnajc3 and Dnajb11 of the Hsp40 family). These genes are required for proper folding, trafficking, or degradation of proteins. Many of these listed genes are functionally coordinated. Specifically, when unfolded (or misfolded) proteins in the ER exceed the capacity of the folding machinery, a redundant set of heat shock proteins (Hsp) initiates the unfolded protein response (UPR). Among the listed genes. Hyou1 carries signals that connect the ER and mitochondria, the main Ca^2+^ storage cell organelles. In this line, calreticulin (Calr), an ER-resident protein, is a Ca^2+^-binding chaperone that acts in neuronal regeneration capacity following injury [47].

**Figure 5.**
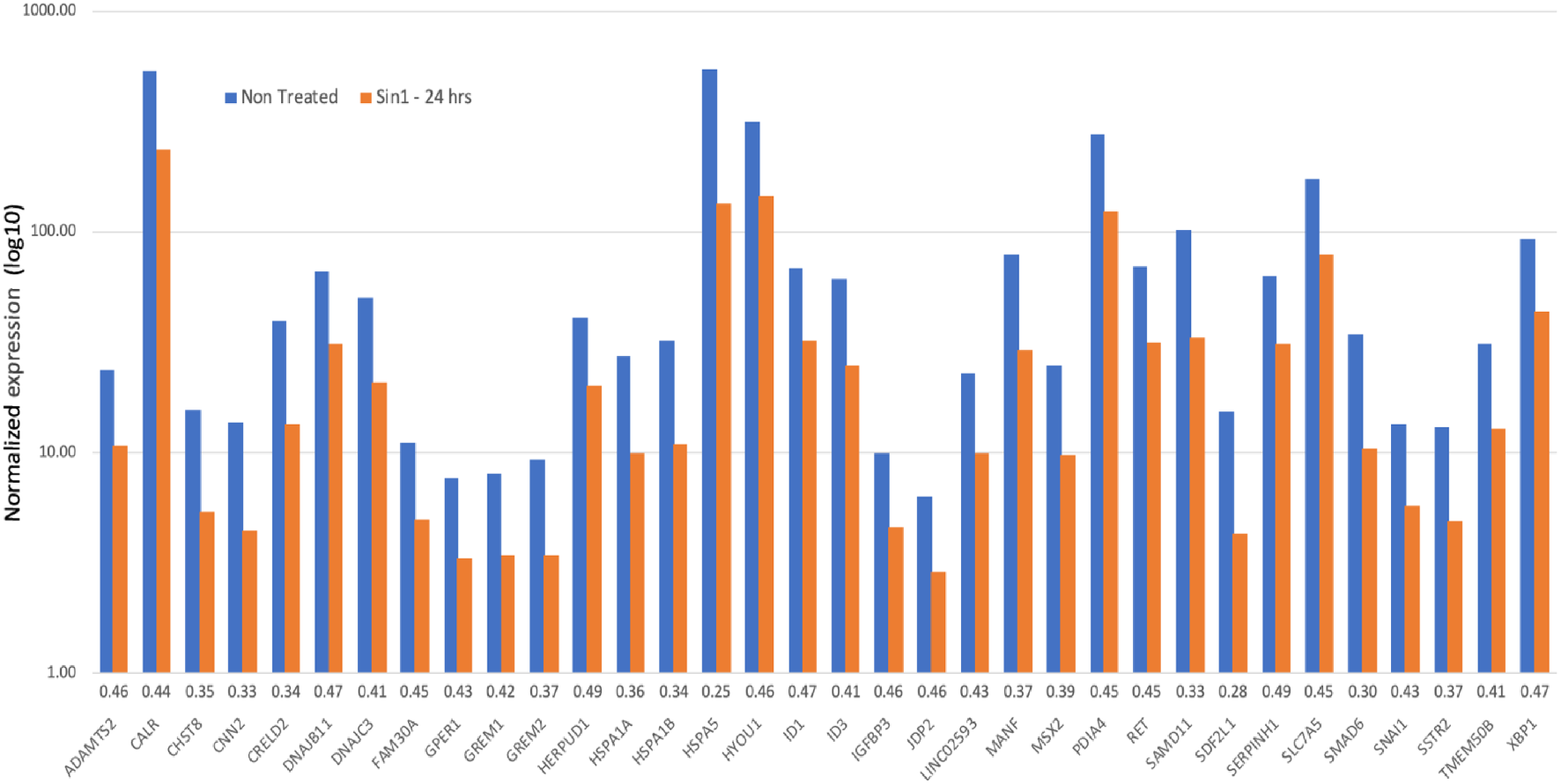
A collection of significant suppressed genes (>2 folds) following exposure to Sin1 with respect to untreated cells (24 hrs). Expression levels are in a logarithmic scale. Genes are sorted alphabetically.

Another gene that is strongly suppressed (0.37) is Manf (Mesencephalic astrocyte-derived neurotrophic factor), an ER-stress regulated protein that exhibits cytoprotective by regulating ER homeostasis [48]. We conclude that the coordinate suppression of the ER maintenance machinery reflects the initial elevation in cell oxidation that potentially suppresses reduced ER protein production in SH-SY5Y cells. We postulate that following oxidative stress induced by Sin1, suppression in the ER misfolding response reflects cells’ adaptation (**Figure 5**).

**Table 2** lists the significantly induced genes (>2-folds). A comparison of induced versus suppressed genes showed that the absolute expression levels were significantly lower (TMM of 11.7 versus 78.8, respectively; p-value = 0.019). Pdk1 (pyruvate dehydrogenase kinase 1; 2.23-fold, p-value = 8.5E-92) is among the few highly expressed induced genes. Overexpression of Pdk1 in animal models and cell lines shows some resistance to Αβ peptides and other neurotoxins [49]. Several of the induced genes are associated with neuronal pathology. For example, Erich3 (2.5-fold, p-value = 7.6E-49) plays a role in vesicular trafficking and neurotransmitter actions, and a change in its expression occurs after antidepressant treatment [50].

**Table 2.**
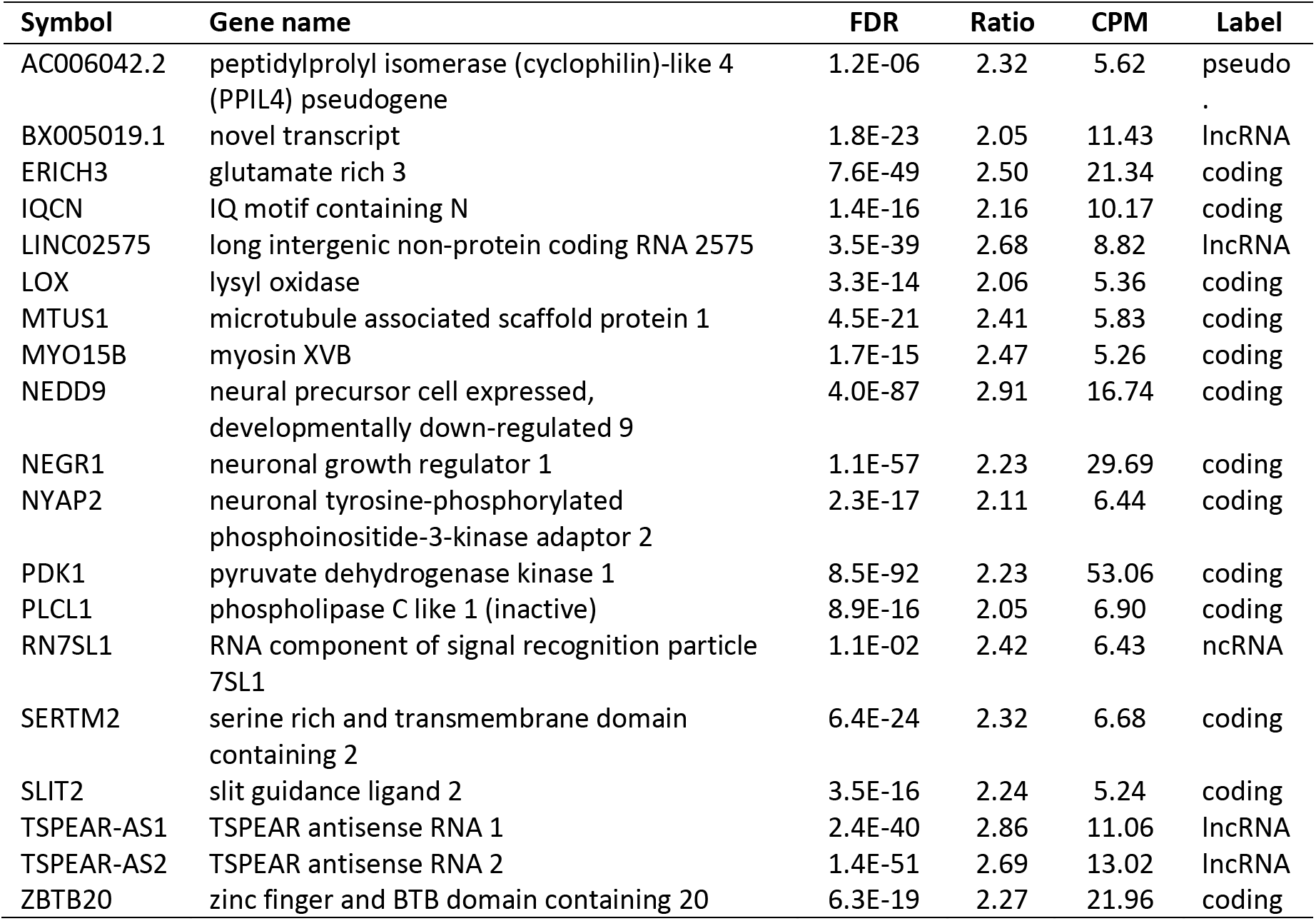
Gene expression induction (>2 folds) in SH-SY5Y exposed to Sin1 (24 hrs) relative to untreated cells

About a third of the significantly induced genes (6 out of 19 genes, >2 fold, >5 TMM) are non-coding RNA, including pseudogenes, antisense, and lncRNA (**Table 2**). Among these ncRNAs, Rn7sl1 (RNA component of signal recognition particle 7SL1) is part of a riboprotein complex that acts in ER translocation. Interestingly, Rn7sl1 was detected in exosomes. Notably, exosomes play a key role in glia-neuronal cell communication [51]. Another notable gene is P2r×7 (Purinergic receptor P2×7; 1.99-fold, p-value FDR = 2.8E-28), an ATP receptor that following its activation, leads to change in Ca2+ homeostasis and an ER-dependent Ca2+ cytotoxicity [52]. A full list of induced genes (n=493) is available in **Supplemental Table S1**.

### Chronic oxidative stress induces a broad range of lncRNAs

Among the Sin1 suppressed genes, 95% are protein-coding genes, and only 3.8% are lncRNAs (**Figure 6**). However, among Sin1 induced genes 83% are protein-coding, and as many as 14% of the induced transcripts are lncRNAs (others are pseudogenes and TEC (transcripts to be experimentally confirmed). Notably, Sin1-induced lncRNAs are mostly low expressing genes. As many as 42% of the listed differential expressed genes are below a threshold of 10 TMM.

**Figure 6.**
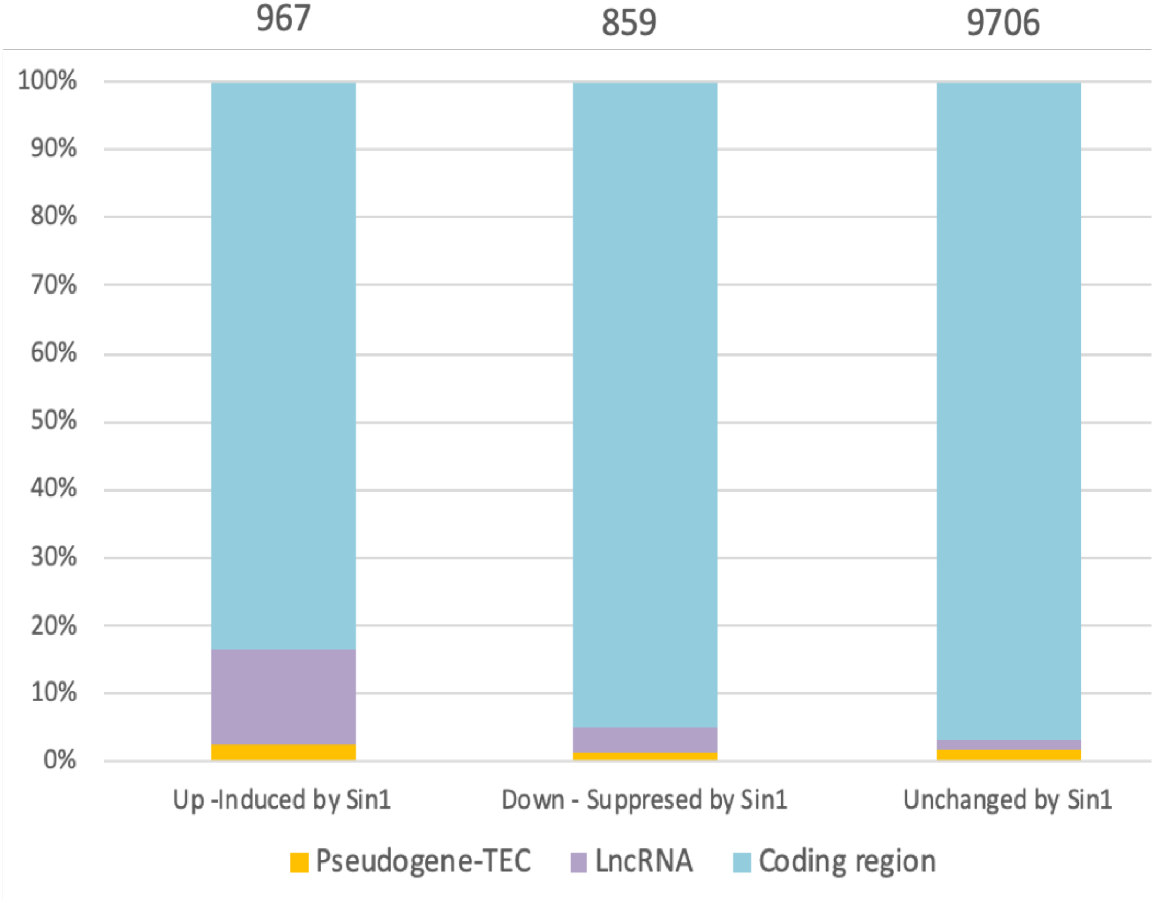
Partitions of genes following Sin1 induction relative to untreated SH-SY5Y cells. Each differential expression group is labelled as coding, lncRNA, and Pseudogene-TEC. The total number of genes is indicated above the histogram bars.

**Table 3** shows the list of 16 lncRNA that is statistically significant and highly expressed (TMM >35) and significantly induced following Sin1 treatment (**Table 3**). The most strongly expressed lncRNAs are Metastasis-associated lung adenocarcinoma transcript 1 (Malat1, p-value FDR = 2.5E-04). Malal1 has also been speculated to be an Nrf2 regulator which is part of a protein complex of the cellular oxidative response and a target that alters oxidative stress through miRNA [53]. Moreover, in recent years, oxidative stress was shown to correlate strongly with lncRNAs expression [54, 55].

**Table 3.** LncRNA genes induced in cells exposed to Sin1 (24 hrs) in comparison to untreated cells.

| lncRNA | lncRNA – Full name | FDR | Folds | TMM<sup>a</sup> |
| --- | --- | --- | --- | --- |
| GATA3-AS1 | GATA3 antisense RNA 1 | 2.3E-08 | 1.32 | 36.5 |
| AL392083.1 | novel transcript, sense intronic to ADARB2 | 1.8E-33 | 1.78 | 37.6 |
| LINC00632 | long intergenic non-protein coding RNA 632 | 5.0E-08 | 1.31 | 40.2 |
| LINC01833 | long intergenic non-protein coding RNA 1833 | 6.8E-14 | 1.32 | 42.0 |
| TTN-AS1 | TTN antisense RNA 1 | 1.2E-06 | 1.26 | 43.0 |
| LINC02268 | long intergenic non-protein coding RNA 2268 | 1.2E-16 | 1.45 | 44.6 |
| GABPB1-IT1 | GABPB1 intronic transcript | 4.6E-15 | 1.35 | 45.6 |
| MIR137HG | MIR137 host gene | 4.5E-23 | 1.47 | 58.7 |
| CCDC144NL-AS1 | CCDC144NL antisense RNA 1 | 1.6E-12 | 1.32 | 67.5 |
| AC093849.2 | novel transcript | 2.9E-33 | 1.46 | 85.0 |
| CASC15 | cancer susceptibility 15 | 1.5E-16 | 1.27 | 85.8 |
| NEAT1 | nuclear paraspeckle assembly transcript 1 | 3.5E-05 | 1.49 | 154.0 |
| MIAT | myocardial infarction associated transcript | 6.3E-21 | 1.35 | 176.9 |
| GABPB1-AS1 | GABPB1 antisense RNA 1 | 1.7E-18 | 1.38 | 280.8 |
| HAND2-AS1 | HAND2 antisense RNA 1 | 6.7E-25 | 1.33 | 671.8 |
| MALAT1 | metastasis associated lung adenocarcinoma transcript 1 | 2.5E-04 | 1.24 | 1963.7 |

<sup>a</sup>TMM, Trimmed mean of M-values

Among the lncRNAs that are highly expressed is the oxidative stress-dependent Neat1. It could reverse the damage caused by superoxide in LPS-treated rodents, and counteract the H_2_O_2_-induced neuronal damage. Neat1 is likely a neuroprotector. A similar function was attributed to Miat. Miat knockdown suppressed viability in stressed cells under hypoxic conditions [56]. Many of the listed lncRNAs genes are involved in competitive endogenous RNAs (ceRNA) used as sponges for miRNAs, others are regulators of Nrf2 function. We conclude that Sin1 induces genes involved in non-classical regulation by lncRNA.

### Ladostigil alters the expression of genes act in dissipating oxidative damage

Incubation of SH-SY5Y cells with ladostigil under basal conditions had a negligible effect on gene expression. Analyzing RNA-seq analysis results at 10 hrs after the introduction of ladostigil relative to untreated cells identified only 7 genes statistically significantly changed genes (total 11,588 genes, **Supplemental Table S1**). Among the downregulated genes is Tmem87a, an endosome-to-TGN retrograde transport (p-value FDR = 5.4E-03) that activates anterograde transport of GPI-anchored and transmembrane proteins. Tmem129 (induced by 24%, a p-value FDR = 0.025) is an ER-resident of protein that is part of the dislocation complex. Several genes that are significantly suppressed but failed to reach the fold change threshold are also ER proteins (e.g., Hspa5, Herpud1) [57]. The abundant mitochondrial NADH-dehydrogenase 6 (MT-ND6) [58] was suppressed by 21%. Oxidative stress by H2O2 led to a diffusion of the MT-ND6 RNA from its matrix location [59]. A dislocation of proteins into the cytosol causes misfolded and topologically impaired ER proteins to be subjected to proteolysis [60]. We conclude that ladostigil per se has a negligible effect on SH-SY5Y cells at a resting state. However, it affected genes associated with the function of the mitochondria and ER.

**Figure 7** shows (by the volcano plot) the effect of ladostigil on gene expression in cells exposed to Sin1. Altogether, only 10 genes (TMM >5; p-value FDR <0.05) resulted in a significant differential expression when compared to Sin1 treated cells (**Supplementary Figure S4**). The most significantly upregulated gene is Clk1 (Cdc-like kinase 1). Clk1 is a multifunctional kinase that phosphorylates serine/arginine (SR) proteins, which affect mRNA splicing. Clk1 was identified as a gene that shows a reduced amount in the hippocampus following chronic stress, while treatment with clomipramine, an antidepressant drug, prevented the stress effect on its expression [61]. As Clk1 was also associated with the pathophysiology of Alzheimer’s disease, it was declared a promising therapeutic target [62]. Inspecting the moderately induced genes (with p-value FDR <0.05) by ladostigil revealed numerous mitochondrial and ER-associated proteins.

**Figure 7.**
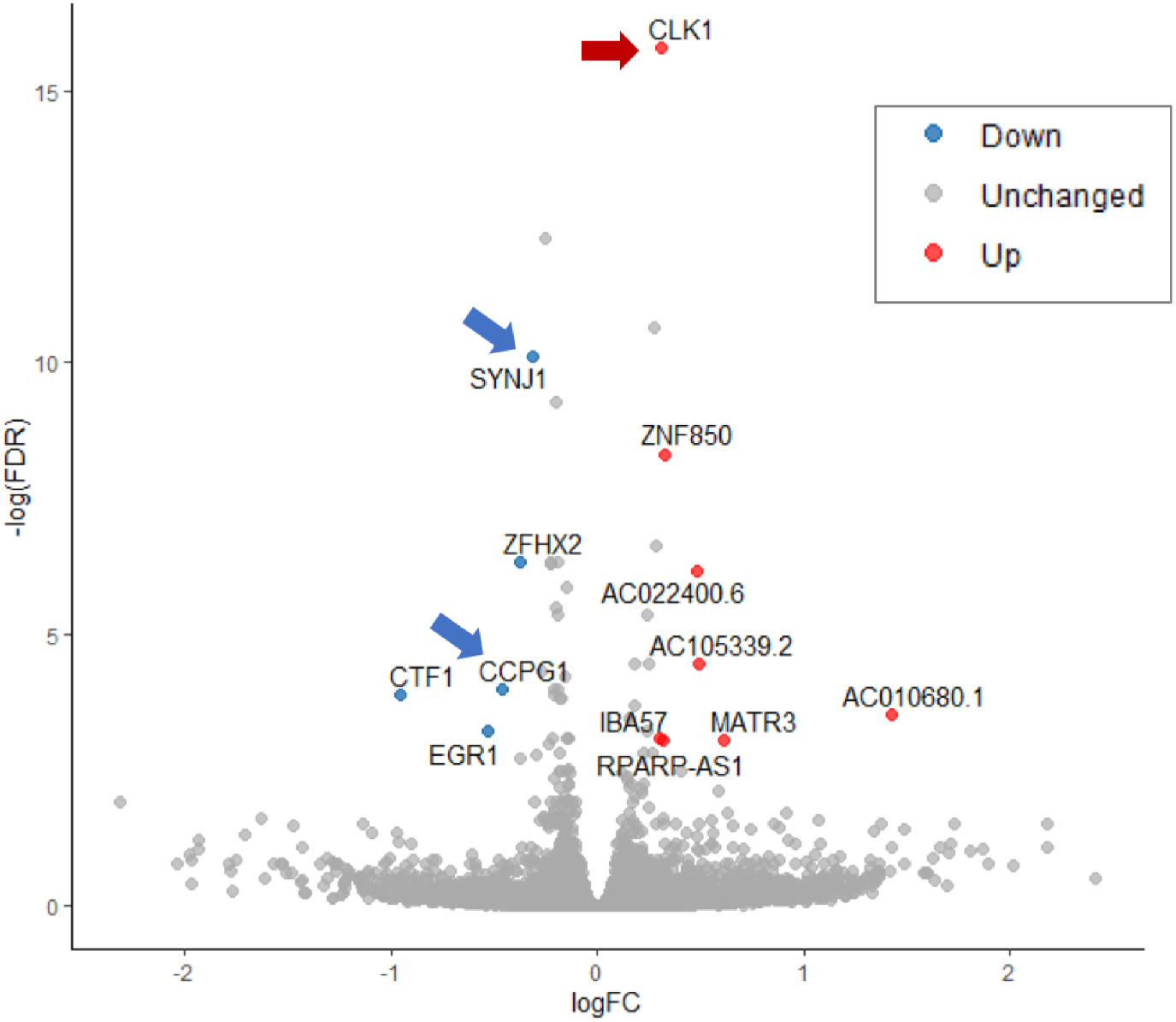
A volcano plot of differential gene expression as a result of pre-incubation (2 hrs) with ladostigil in the presence of Sin1 (24 hrs) compared to cells that were exposed to Sin1 alone. Three biological replicates were used for the RNA-seq analysis. The log FC (fold change) was calculated by edgeR (with FDR < 0.05). Arrows mark the statistically significant genes that function in oxidative and ER stress response.

Among these genes are Iba57 (Iron-sulfur cluster assembly factor Ia57), a protein that is essential for the maturation of mitochondrial iron [4Fe-4S] protein complex [63]. The expression level of Matr3 (Matrin3) increased by 1.5-fold. Mutations in this gene lead to amyotrophic lateral sclerosis (ALS) and frontotemporal dementia (FTD). Functional impairment of Matr3 and its localization in the nucleus induced proteotoxicity associated with these diseases [64]. Txnip (Thioredoxin interacting protein) is among the gene that was significantly differentially expressed (albeit below that default 1.33-fold threshold). It was proposed a linker of ER stress and oxidative stress in the formation of inflammasome [65]. Similarly, Atg14 (Autophagy related 14), a component of the autophagy-specific membrane fusion [66] was slightly increased by ladostigil (p-value FDR = 1.2E-03). The rest of the genes are ncRNA transcriptional regulation genes (with pseudogene, retained introns, and a super-enhancer). **Supplementary Table S2** lists the differentially expressed genes along with their statistical significance.

Synaptojanin 1 (Synj1, p-value FDR = 2.1E-05) is among the very few downregulated genes. The role of Synj1 is to facilitate the recycling of synaptic vesicles in neurons. Synj1 role in regulating autophagy was reported based on Synj1-deficient mice that were associated with an impairment in age-dependent autophagy in multiple brain regions [67]. Moreover, the reduction of Synj1 accelerated the clearance of the toxic Aβ peptide and consequently attenuated cognitive decline [68]. The other ladostigil suppressed gene is Ccpg1 (cell-cycle progression gene 1), an ER-resident protein that participates in autophagy of the ER (denoted ER-phagy) and membrane dynamics. In numerous cells, the Ccpg1 gene was induced by the UPR, thus directly links ER stress to ER-phagy [69]. We postulate that the Sin1 partially prevents the UPR response (e.g., XBP1, **Figure 4**), and thus ER-phagy is further diminished by ladostigil.

## Discussion

A significant reduction in the age-dependent decline of spatial memory and learning was observed following a long-term treatment of rats by ladostigil (1 mg/kg/day; for 6 months). The anti-inflammatory activity mediated by microglia underlies the neuroprotective action of ladostigil [22, 26]. Results from RNA-seq analysis of aged rat brains revealed substantial alterations in gene expression in different brain areas as a consequence of long-term ladostigil treatment [21] **(Figure 1)**. However, the complexity of the cell composition in the brain (e.g., microglia, neurons, astrocytes) masked cell-specific effects.

Cells that are strongly dependent on mitochondrial function (e.g., neurons, muscle) and protein secretion (e.g., neurons, microglia, neuroendocrine) are suitable to investigate stress-dependent mechanisms. In this study, we used undifferentiated SH-SY5Y cells as a model for CNS neurons. The SH-SY5Y cells exhibit neuron-like [32], primarily the inter-connection between ROS (mitochondrial and cytosol) and ER proteostasis. Numerous rare diseases and common chronic conditions highlighted such interplay between oxidative and ER stress [70].

The effect of ladostigil is manifested through multiple pathways (e.g., adhesion, neurogenesis, calcium homeostasis) that lead to a reduction in microglia activation [21]. Studying the molecular signature underlying ladostigil’s action in the brains of aging rats showed a reduction in crucial phosphorylation sites that drive Nf-κB translocation to the nucleus [26]. Nf-κB is a bridge between pro-inflammatory cytokine expression and neurogenesis [71, 72]. Therefore, it is an attractive therapeutic route for neurodegenerative diseases [73].

In this study, we tested the impact on gene expression 24 hrs after stress induction. We showed that under acute exposure to H_2_O_2_, SH-SY5Y cells have a substantial capacity to overcome the accumulated damage (**Figure 2**). The pathology of brain aging is associated with an imbalance between the antioxidant response mechanism and internal ROS generation [74]. We showed that under irreversible damage by a high concentration of oxygen peroxide, ladostigil fails to rescue cells from apoptotic death (**Figure 2**). Notably, in cultured cells, H_2_O_2_ can only act for a few minutes [75]. In general, excess oxidants may compromise cell function by modifying proteins, lipids, and cell energetics, ultimately leading to cell death. Our results emphasize the difference between acute, short-lived stress (e.g., by H_2_O_2_) and a long-lived stressor (e.g., Sin1). Sin1 is a source of reactive peroxide that causes lipid peroxidation and mitochondrial dysfunction, and its impact on cells lasts for hours [76, 77]. We Showed that priming cells by ladostigil before Sin1 enables cells to better cope with chronic stress (**Figure 3**).

Suppression of genes involved in ER, chaperons, and quality control **(Figure 4**) signifies a coordinated response of Sin1-exposed cells (50 μM, 24 hrs). Inspection of the expression profile of these Sin1 exposures revealed a link between the redox state and protein folding homeostasis (**Figure 5**). At the organism level, failure in maintaining a healthy redox state triggers pathological conditions that eventually lead to diseases (e.g., neurodegenerative, cardiovascular, metabolic disorders). In the aging brain, it is mostly the astrocytes and activated microglia that secrete extracellular cytokines that lead to accelerated ER malfunction. Under mild ER stress, the UPR system is turned on, in parallel to the shut-down of protein production. An increased level of inflammation (**Figure 1**) leads to dysfunction of the UPR pathways [78]. In SH-SY5Y cells, ladostigil altered just a handful of genes that act in the ER quality control and ER-phagy (**Figure 7**). We propose that protein overload (due to a suppressed capacity to resolve misfolding and aggregates) can lead to mitochondrial function disturbance. Probably, the toxic accumulation of ROS and nitroperoxides in cells affects the integrity and function of organelles which trigger autophagy (ER-phagy and Mitophagy) [69].

The enrichment in ncRNAs following Sin1 chronic exposure is an unexpected observation (**Figure 6, Table 2**). However, in Parkinson’s disease the introduction of synthetic α-synuclein altered lncRNAs expression profile [79]. The expression of ncRNAs signifies the ability of cells to cope with several types of stress conditions (metabolic stress, oxidative stress, genotoxic stress) [80]. An attractive possibility concerns the role of ncRNA in exosomal communication in aging and inflammation [51].

In summary, we characterized SH-SY5Y as an attractive model for exposing the impact of mild but chronic stress on neuron-like cells. We argue that the molecular response is likely to be cell-type specific. Thus, cells with low antioxidative activity are prone to damage by external insults [81]. Neurons are exceptionally sensitive to nitrogen and oxygen-based damage due to their dependence on the respiratory chain [82]. We propose that the beneficial effect of ladostigil is on a failure of neurons in the aging brain to maintain effective sorting, trafficking, and secretion. Under chronic oxidative stress, the redox homeostasis is disturbed, and consequently, the protein folding and ER function become fragile. Ladostigil affected a small set of genes including numerous lncRNAs and genes that regulate ER-autophagy and endocytosis in SH-SY5Y cells. We conclude that regulating ER stress and attenuating oxidative stress by ladostigil may reduce the pathophysiology outcomes of aging, brain injury, and neurodegenerative diseases.

## Abbreviations

ALS: amyotrophic lateral sclerosis
CNS: central nerve system
DE: differentially expressed
ER: endoplasmic reticulum
FACS: fluorescence-activated cell sorting
FC: fold change
FCS: fetal calf serum
FDR: false discovery rate
FTD: frontotemporal dementia
GO: gene ontology
LPS: lipopolysaccharide
ROS: reactive oxygen species
TMM: trimmed mean of M-values

## Acknowledgments

We would like to thank Marta Weinstock-Rosin (School of Pharmacy, The Hebrew University of Jerusalem) for providing us with ladostigil, and for helpful discussions throughout this research. We thank Dana Reichman (Life Science Institute, (School of Pharmacy, The Hebrew University of Jerusalem) for providing us the GFP-redox-sensors plasmids.

## Availability of data and material

Normalized data is available in supplementary Table S1 and DE (differential expressed) is available in Supplemental Table S2. The raw data is shared in ArrayExpress accession E-MTAB-10817.

## Ethics approval

All participants have completed CITI program ethical course for good practice in human and animal research. The study is limited to research on established cell line.

