## Supplemental Materials for "Ladostigil attenuates the oxidative and ER stress in human neuroblast-like SH-SY5Y cells"

**Supplemental Table S1**: Normalized data (by TMM) for all expressed genes >5 TMM

**Supplemental Table S2**: Differentially expressed genes and their statistical significance.

### Supplementary Figures

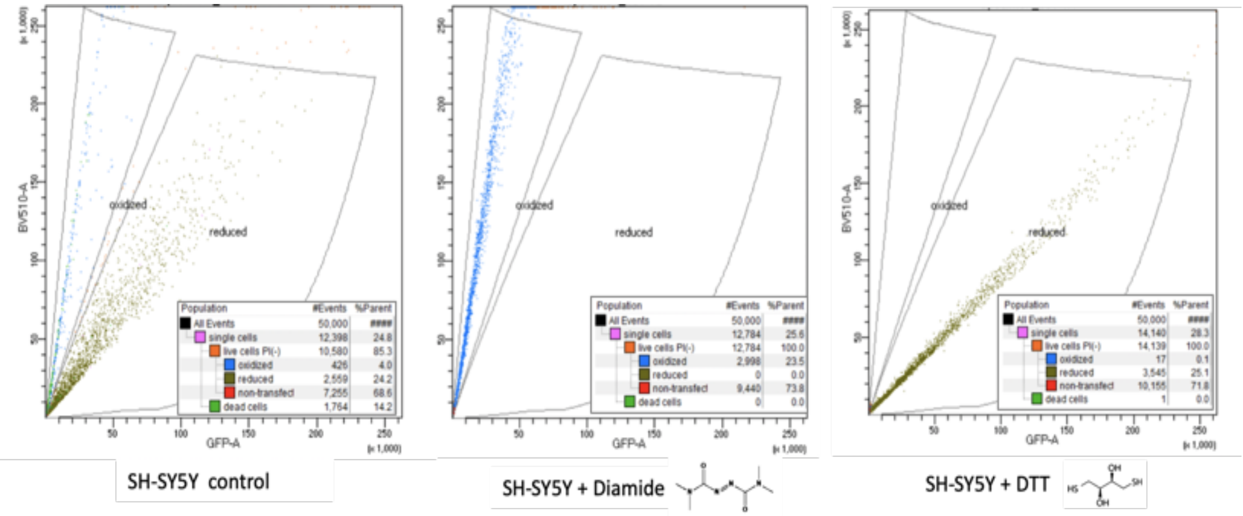


**Figure S1.** Calibration of redox level in cells**.** Cells were transfected with redox-measured plasmids with GFP. The transfected cells were treated after 24 hrs. The FACS gating of the extreme conditions are used as internal calibration. The FACS monitored cells after incubation with diamide which shifted all living cells toward the oxidized state (middle), and with DTT, that shifted cells to a maximal degree of reduced and oxidized states (right). The control of untreated cells (left) shows that both reduced and oxidized states can be quantified. 50,000 cells were quantified for each condition.

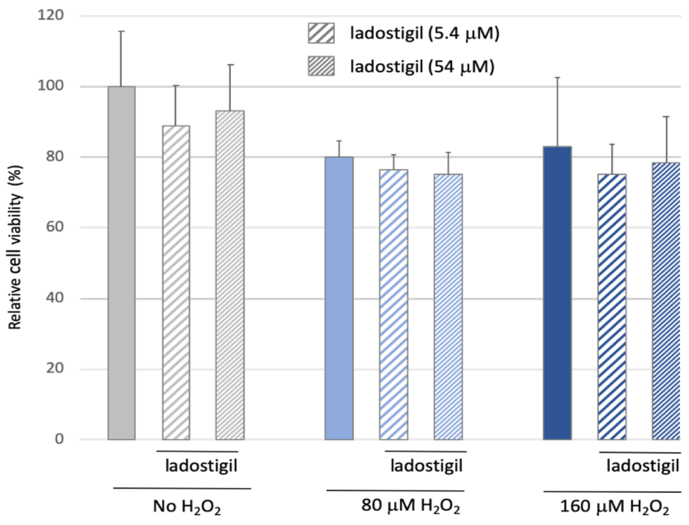


**Figure S2.** SH-SY5Y cells under acute short-term oxidation stress and ladogtigil treatment. Monitoring the impact on cell viability in the presence of two concentrations of ladogtigil and oxygen peroxide. Each histogram bar represents the % relative to naïve untreated cells measured in 8 wells. Ladostigil was shown to be non-toxic over a wide range of concentrations.

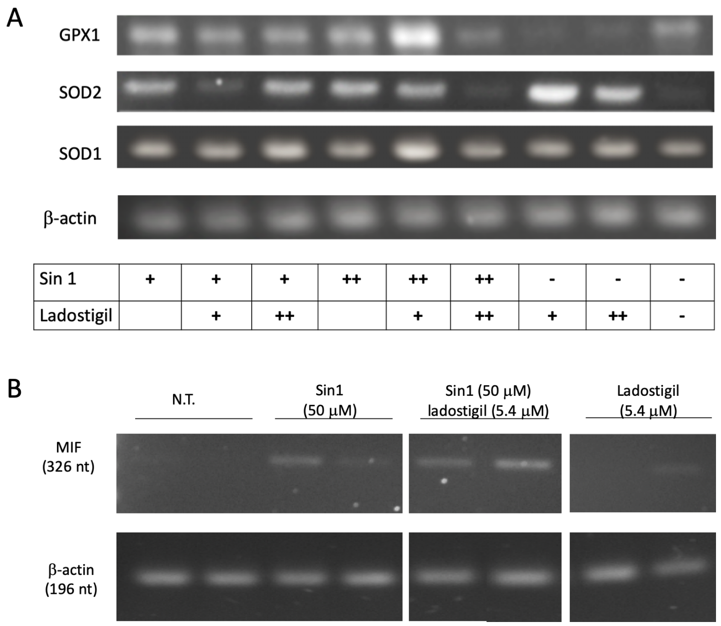


**Figure S3.** Gene expression of major ROS detoxification enzymes in SH-SY5Y cells under Sin1 and ladostigil. **(A)** The results of RT-PCR for the antioxidant superoxide dismutase enzymes (Sod1 and Sod2). Cytoplasmic Sod1, was slightly induced by Sin1, while Sod2, a mitochondrial resident enzyme, was induced >2-fold from its basal level. Ladostigil at 5.4 μM (but not 54 μM), reduced Sod2 to its basal level. A similar pattern was recorded for Glutathione peroxidase 1 (Gpx1), a ubiquitous enzyme that is located in the cytosol, mitochondria, and peroxisomes. The low and high concentrations of Sin1 (50 and 100 mM) and ladostigil (5.4 μM and 54 μM) are marked + and ++, respectively. **(B)** Mif (Macrophage migration inhibitory factor) is slightly induced by Sin1 but ladogligil induces its expression even further. Each condition was tested on separated biological samples from RNA extracted from two cell cultures.

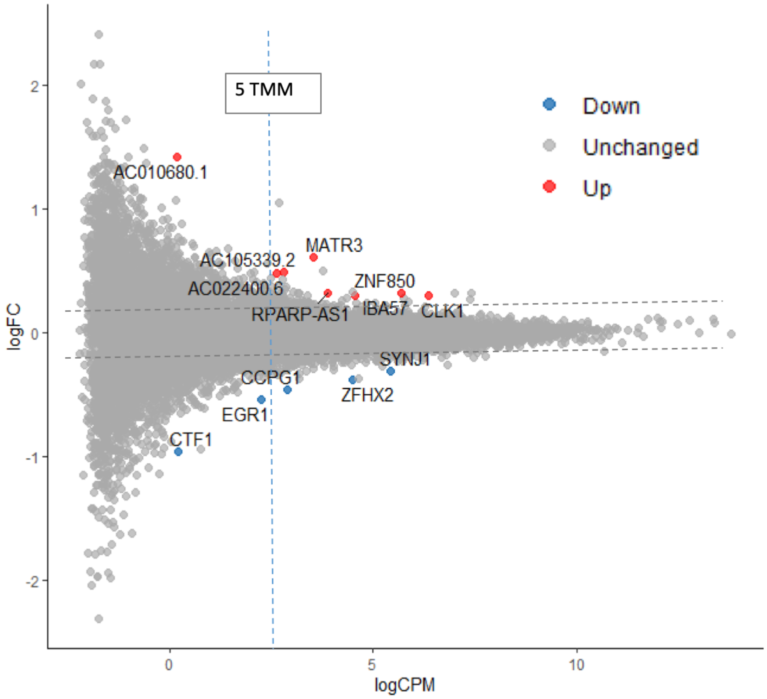


**Figure S4.** A plot of differential gene expression as a result of pre-incubation (2 hrs) with ladostigil in the presence of Sin1 (24 hrs) compared to cells that were exposed to Sin1 alone. Three biological replicates were used for the RNA-seq analysis. The log FC (fold change) was calculated by edgeR (with FDR < 0.05). LogCPM shows the absolute expression level (TMM). The threshold of TMM=5 and FC are marked by dashed lines.
